# Genome Mining of *Salinispora* Uncovers a Repertoire of Secondary Metabolites and Carbohydrate-Active Enzymes

**DOI:** 10.1101/2023.09.29.560222

**Authors:** Md. Saddam Hossain, Md. Tarikul Islam, Md. Hadisur Rahman, Abdullah Al Zubaer, Abu Hashem

## Abstract

Natural products derived from microorganisms are a significant part of the pharmaceutical industry’s arsenal today. These efforts have increasingly centred on marine microbes, which have produced a variety of intriguing novel chemicals. Marine-derived strains are likely candidates for natural product discovery since actinomycetes are a significant source of microbially produced natural compounds. The availability of microbial genome sequencing has revealed a number of natural products (NPs) that have not yet been thoroughly investigated. The genome mining potential of the genus *Salinispora*, which has potential for the production of secondary metabolites of clinical importance, remains unexplored. In this study, we have demonstrated a thorough comprehension of the genetic characteristics, biosynthetic gene clusters (BGCs), and genetic clusters for carbohydrate-active enzymes found in the *Salinispora* genomes. In our analysis, we have identified 686 BGCs, among which 3.79% are identical to the BGCs that produce ketomemicin B3/ketomemicin B4, salinilactam, lymphostin/neolymphostinol B/lymphostinol/neolymphostin B, benzastatin, and thiolactomycin. We have found 19.5% of the BGCs, which are only present in the genomes of *Salinispora* and may open the door to the development of novel drugs. We have also found a total of 3604 genes in 62 various categories of the six major carbohydrate-active enzymes. The development of new-generation bioactive compounds that are significant for biotechnology relies heavily on the scientific insights provided by genome mining in this genus.

## Introduction

Actinobacteria are a rich source of secondary metabolites that have clinical significance, particularly as anesthetics, immunosuppressants, antibacterial, antifungal, antiviral, and anticancer agents [1]. In addition, a variety of food additives and crop protection substances come from natural product (NP) sources of bacteria, fungi and plants [2]. Approximately 65% of currently used clinical medications and more than 50% of Food and Drug Administration (FDA)-approved drugs are NP-inspired. These compounds originate from numerous chemical classes, including polyketides (PKs) [3], nonribosomal peptides (NRPs) [4], ribosomally synthesized and post-translationally modified peptides (RiPPs) [5], saccharides [6], alkaloids [7], and terpenoids [8], which collectively cover an astounding variety of chemical scaffolds.

Finding new drugs with clinical relevance is crucial because rapidly emerging multi-drug resistant (MDR), the gram-negative *Enterococcus faecium*, *Staphylococcus aureus*, *Klebsiella pneumoniae*, *Acinetobacter baumannii*, *Pseudomonas aeruginosa*, and *Enterobacter spp*. (“ESKAPE” bacteria), and extensively drug-resistant (XDR) pathogens are posing an increasing threat to several potentially fatal diseases like cancer, Alzheimer’s, cirrhosis, tuberculosis, pneumonia, influenza, etc [9,10]. The primary method for discovering biologically and therapeutically significant natural products is the extraction of culture filtrates, but this approach falls short because a large number of chemicals remain unexplored [11]. Intriguingly, the majority of natural products originating from microorganisms (and some from plants) are produced by metabolic pathways encoded by chromosomally nearby genes called biosynthetic gene clusters (BGCs), which encode the enzymes, regulatory proteins and transporters that are necessary to produce, process and export secondary metabolites. Importantly, these features make it possible to mine the microbial genome for metabolites using computational methodologies and facilitate the identification of cryptic BGCs that were unable to produce desirable end products under laboratory conditions [12,13]. Effective investigation of the vast range of compounds produced by organisms throughout the tree of life is eventually made possible by in-silico analysis of the bacterial draft or complete genome sequence data. Additionally, this method has shown promise for discovering new medications because it appears to cut production costs and time while also reducing the need to repeatedly isolate the same chemical components [12].

The quest for new secondary metabolites has shifted its focus to actinomycetes generated in marine environments [14]. The strains of *Salinispora* which were first reported in 1989 from a salty environment (marine habitat) are aerobic, gram-positive, non-acid-fast actinomycetes that produce highly branching substrate hyphae that form single or multiple smooth-surfaced spores [15,16]. It only has 9 type species that have been identified internationally as of this writing (https://lpsn.dsmz.de/search?word=salinispora). This genus, which is widely dispersed in tropical and subtropical ocean sediments, serves as an unusually rich source of structurally varied secondary metabolites that are synthesized according to species-specific patterns [17]. Many new natural products, such as salinosporamide A for the treatment of cancer, sporolides A and B, etc. from *Salinispora* have been discovered and characterized [18,19]. The NCBI Reference Sequence (RefSeq) collection provides a comprehensive, integrated, non-redundant, well-annotated set of sequences, including genomic DNA, transcripts, and proteins. We have scanned all publicly available reference genome sequences of species in the *Salinispora* genus using a variety of bioinformatics tools to examine any potential biosynthetic clusters, putative products of these BGCs, evolutionary relationships of PKS KS and NRPS C domains, and CAZymes (carbohydrate-active enzymes) that may be present. A significant number of gene clusters involved in the manufacture of novel potential chemicals of medical and agricultural significance have been found by screening 27 *Salinispora* genomes.

## Materials and Method

### Retrieval of Salinispora WGSs

All *Salinispora* genomes from the NCBI Refseq genome dataset (https://www.ncbi.nlm.nih.gov/data-hub/genome/) were downloaded (last browsed in March 2023). We have found a total of 30 genomes at the contig, scaffold and complete genome level and we downloaded 27 genomes and avoided 3 genomes as they were suppressed in the NCBI database, which means they failed to pass the genome assessment. We also searched the EzBioCloud (https://www.ezbiocloud.net/) database and found 91 genomes, but when we searched the respective GenBank accession number in the NCBI database, we found them suppressed, excluding 27 genomes that were downloaded from the NCBI Refseq genome dataset. The goal was to filter out poorly assembled genomes which would result in a significant underestimation of the secondary metabolite prediction.

### A Comparative Study of the Available Genomes

The average nucleotide identity (ANI) value was calculated with an improved ANI algorithm, called OrthoANI using EzBioCloud (Yoon et al., 2017), and JSpeciesWS (https://jspecies.ribohost.com/jspeciesws/#analyse) was also used to check the genetic diversity among genomes. Phylogenomic analysis based on Genome BLAST Distance Phylogeny (GBDP) was performed using TYGS (Type strain genome server) to determine the relationship between all the *Salinispora* species(Meier-Kolthoff & Göker, 2019) and interactive tree of life (iToL version 6.7.5) was used to visualize the phylogenetic tree. To analyze genomics data like open reading frames (ORFs), GC content, and orthologous groups, each genome sequence was automatically executed on the M1CR0B1AL1Z3R web server (https://microbializer.tau.ac.il/index.html) with the default parameters (Avram et al., 2019). To build a circular map of each genome, all genomes were aligned and compared using the BLAST Ring Image Generator (BRIG) (Alikhan et al., 2011). Using cogclassifier 1.0.5, a Python-based package, all the annotated protein sequences were examined to analyze functional classifications of clusters of orthologous groups (COGs).

### Identification of Gene Clusters Potentially Encoding Secondary Metabolites

To explore the known and unknown secondary metabolite biosynthetic potential all throughout the *Salinispora* genomes, we used the antibiotics and secondary metabolite analysis shell (antiSMASH v7.0.0) bacterial version as web server which makes it easy to find, annotate, and research secondary metabolite biosynthesis gene clusters (Blin et al., 2023). Prediction Informatics for Secondary Metabolomes (PRISM version 4.4.5) was employed to analyze secondary metabolite structure and biological activity in a complete manner, and the BAGEL4 tool was used to comprehensively mine Ribosomally synthesized and post-translationally modified peptides (RiPPs) and bacteriocin (Skinnider et al., 2020; van Heel et al., 2018). antiSMASH job IDs were submitted to the biosynthetic gene cluster family database (BiG-FAM) for gene cluster family (GCF) explorations and representations of annotated BGCs (Kautsar, Blin, et al., 2021). The biosynthetic gene similarity clustering and prospecting engine (BiG-SCAPE) (Navarro-Muñoz et al., 2020) with a distance cut-off score of 0.3, and “mix” and “singletons_included” parameters was used to generate a sequence-based similarity network of the detected clusters, and results were visualized as a network using Cytoscape 3.9.1(Shannon et al., 2003).

### Analysis of Natural Product Domains

Ketosynthase (KS) and condensation (C) domains were detected and classified from the *Salinispora* genomes using the NaPDoS2 web server (https://npdomainseeker.sdsc.edu/napdos2/) version 13b with default parameters.

### Prediction of Carbohydrate Active Enzymes (CAZymes)

dbCAN3 (automated carbohydrate-active enzyme and substrate annotation) web server, which annotates proteins using DIAMOND, HMMER via CAZy, dbCAN, and dbCAN-sub, respectively was used to identify genes that code carbohydrate active enzymes (CAZymes) (Zheng et al., 2023). We kept the enzymes, that hit at least two tools out of three. Additionally, CAZyme gene clusters (CGCs) and the signal peptides were detected by dbCAN3 in all the *Salinispora* genomes.

## Results

### Salinispora Genome’s General Characteristics

Out of 27 gonomes, only two strains are at the complete level (*Salinispora tropica* CNB-440 and *Salinispora arenicola* DSM 44819), and others are at the scaffold and contig levels, which means they are draft genomes. The ranges of genome size, GC (%) content, and total number of genes are from 5.03 MB to 5.9 MB, 69.2% to 70.1%, and 4740 to 5479, respectively. *Salinispora mooreana* CNT-150 and *Salinispora arenicola* CNS673 each have the smallest and largest genomes, respectively. For all the studied genomes, completeness is above 97% and contamination is below 2%, as estimated by CheckM (**Table 1 and Table S1**).

**Table 1.**
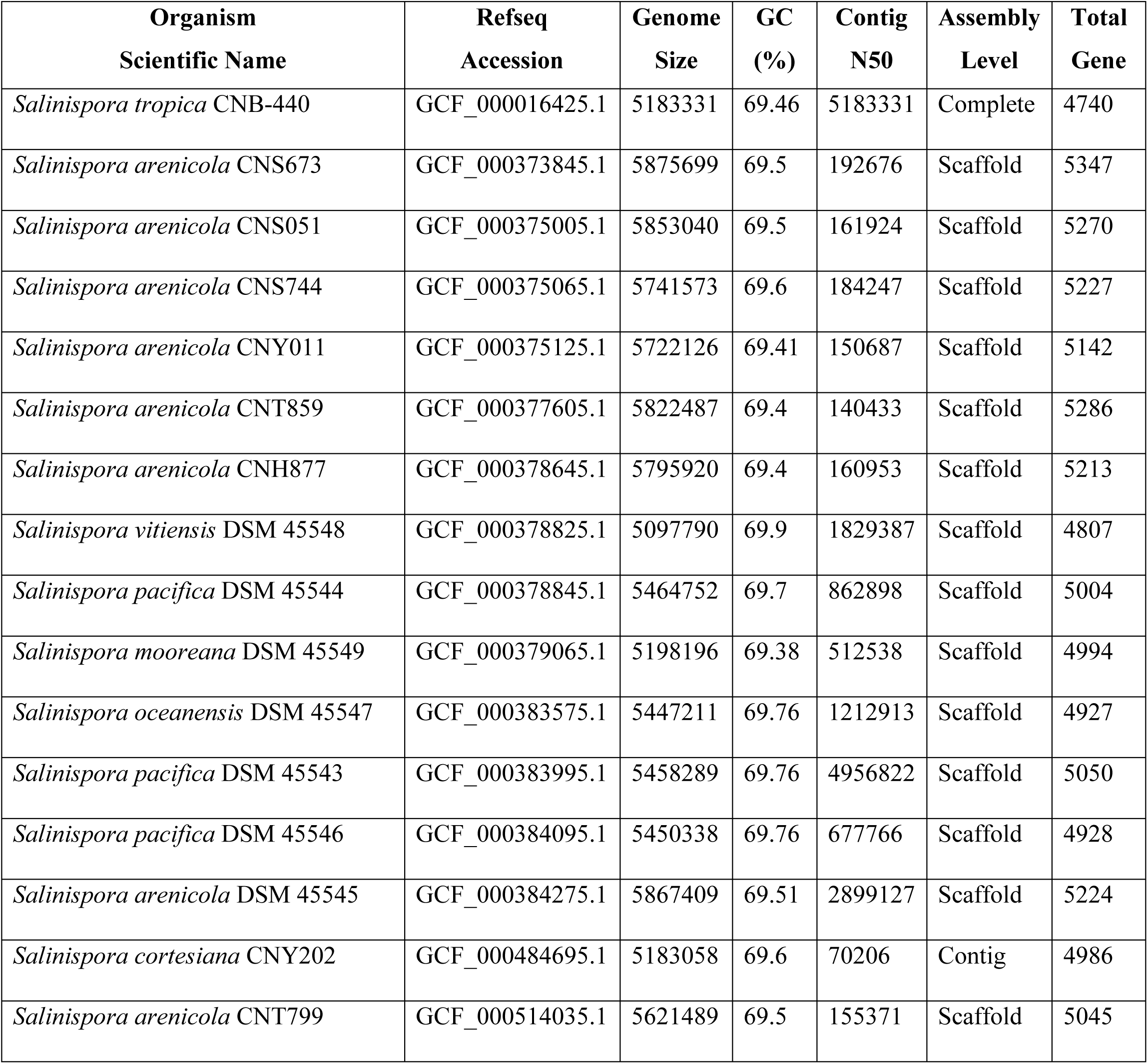
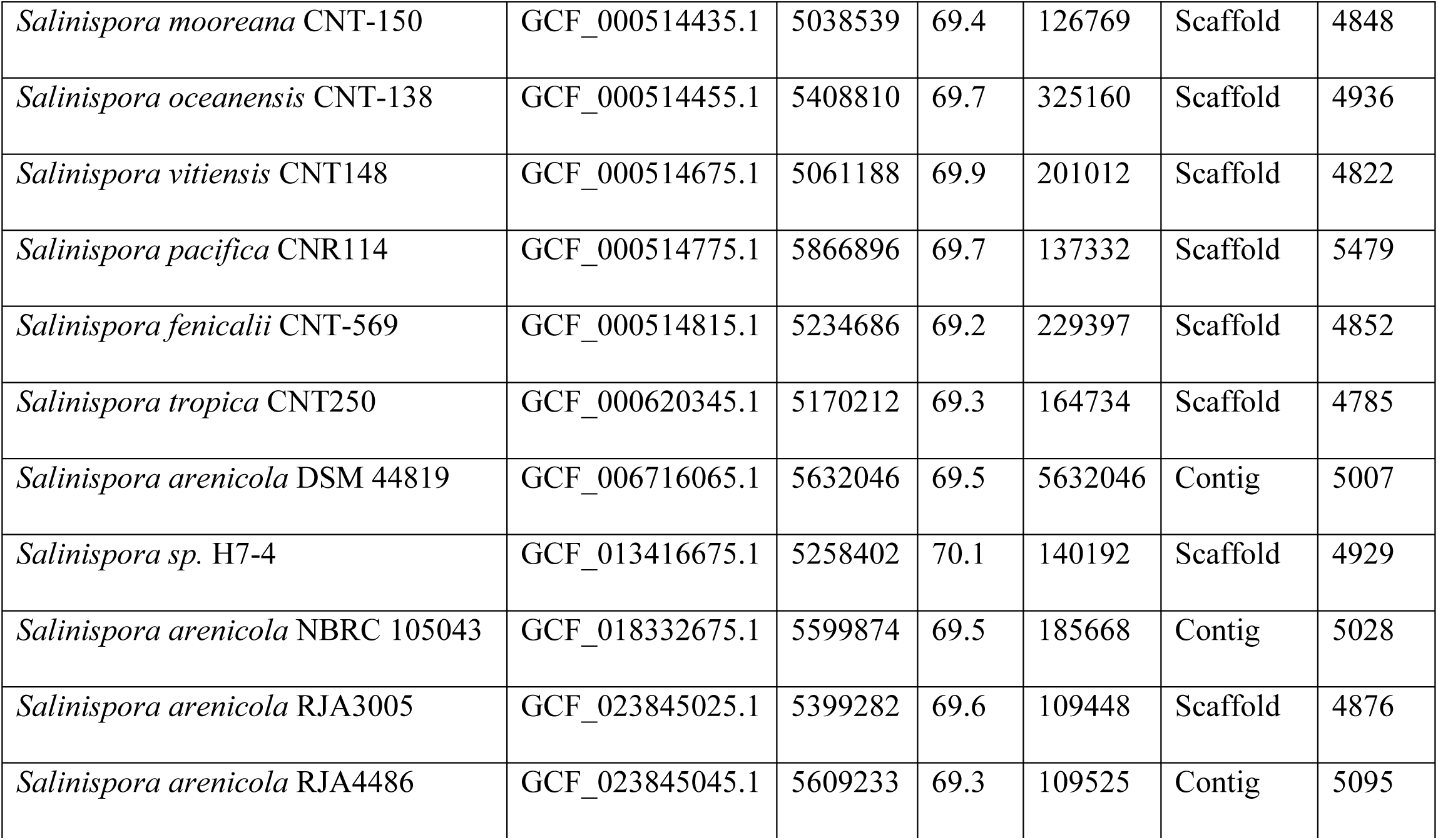
Genomic characteristics of the studied genomes of *Salinispora* species.

### Whole Genome Comparison of Salinispora species

Out of the 27 genomes that were looked at, 10511 orthologous genes were found, and 2603 of them were found to be part of the core proteome. The core proteome is made up of genes that are found in all organisms and has a length of 906911 nucleotides. The number of ORFs present in *Salinispora* genomes ranges from 4607 to 5431 (**Figures S1, S2 and S3**).

**Figure 1.**
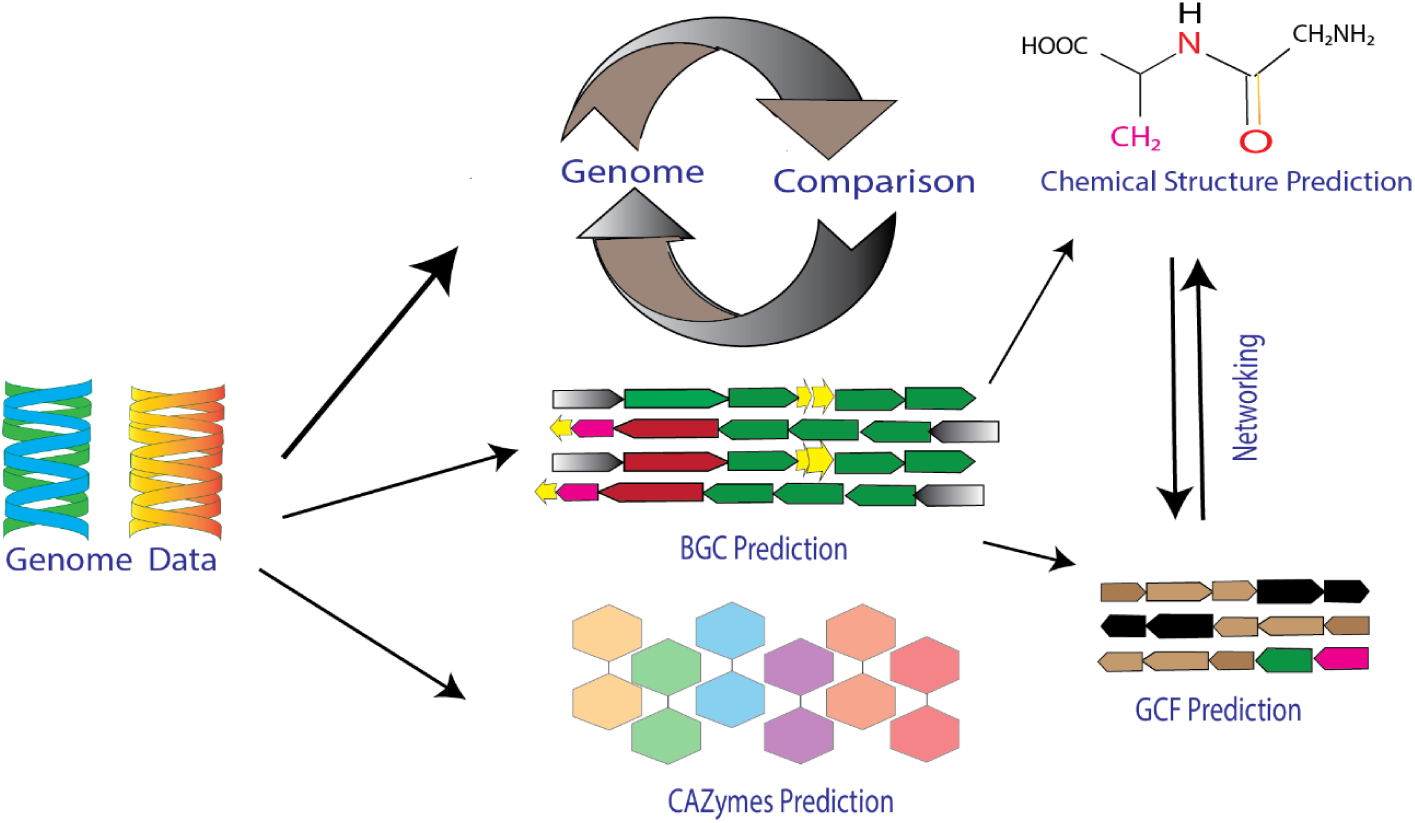
A graphical representation of the workflow.

There are a total of 26 functional categories in the COG system (Galperin et al., 2018). In our study we observed 23 functional categories of COGs protein including Transcription (K), Amino acid transport and metabolism (E), Coenzyme transport and metabolism (H), Translation, ribosomal structure and biogenesis (J), Lipid transport and metabolism (I), Carbohydrate transport and metabolism (G), General function prediction only (R), Energy production and conversion (C), Signal transduction mechanisms (T), Cell wall/membrane/envelope biogenesis (M), Posttranslational modification, protein turnover, chaperones (O), Inorganic ion transport and metabolism (P), Replication, recombination and repair (L), Secondary metabolites biosynthesis, transport and catabolism (Q), Function unknown (S), Nucleotide transport and metabolism (F), Defense mechanisms (V), Cell cycle control, cell division, chromosome partitioning (D), Mobilome: prophages, transposons (X), Intracellular trafficking, secretion, and vesicular transport(U), Cell motility (N), RNA processing and modification (A), and Extracellular structures (W) which are grouped into four main classes namely (1) information storage and processing, (2) cellular processing and signaling, (3) metabolism, (4) poorly characterized (Figure 2).

**Figure 2.**
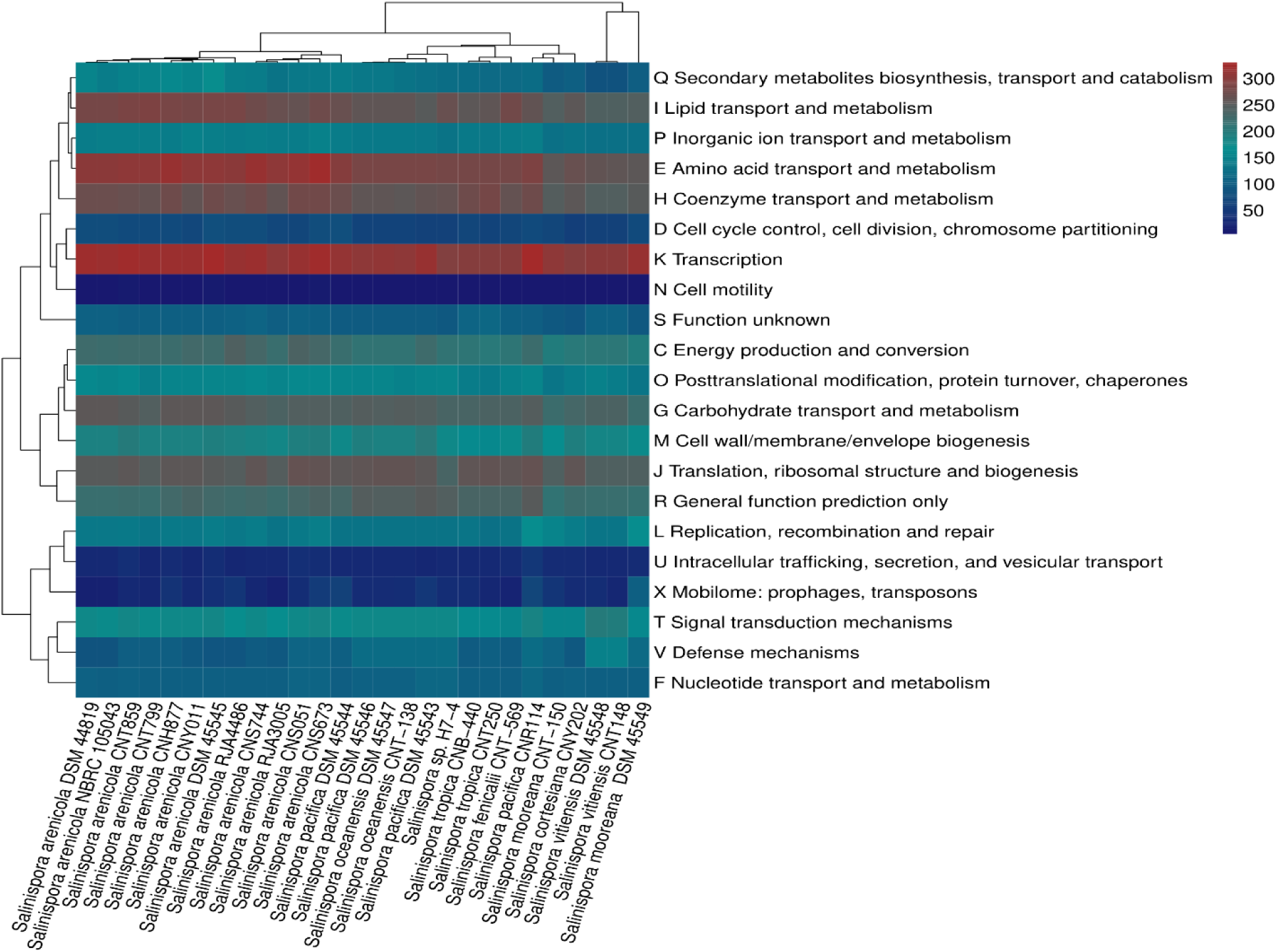
Abundance of Clusters of Orthologous Groups (COGs) in *Salinispora* genomes: Blue and red represents higher and lower abundances respectively.

COG proteins are mainly engaged in Transcription (K), Amino acid transport and metabolism (E), Coenzyme transport and metabolism (H), Translation, ribosomal structure and biogenesis (J), and Lipid transport and metabolism (I). The complete genome of *Salinispora tropica* CNB-440 was used to compare it against other genomes using a circular map produced using BRIG version 0.95.

The map displayed gaps or zones of low similarity, which suggest variances in many regions (Figure 3).

**Figure 3.**
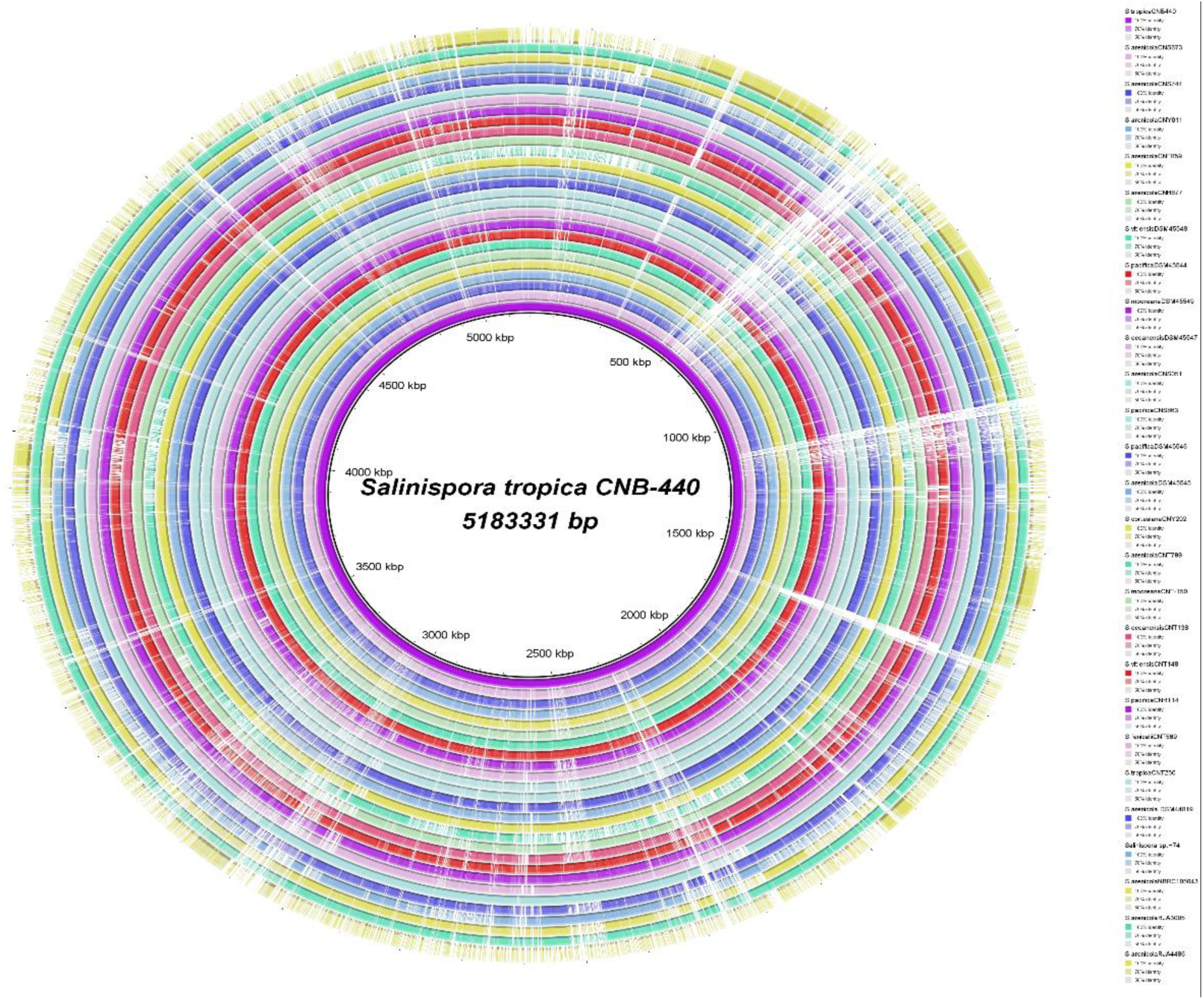
A circular genome map of the entire *Salinispora tropica* CNB-440 as well as comparative genome maps of the other *Salinispora* genome sequences that are currently available.

The Genome-to-Genome Distance Calculator (GGDC) approach is used based on Average Nucleotide Identity (ANI), Digital DNA-DNA hybridization (dDDH) values and differences in percent genomic G+C content (cut off at 96%, 70% and 0.1%, respectively) to separate an organism at the species and sub-species level. Except for *Salinispora oceanensis* DSM 45547 and *Salinispora pacifica* DSM 45546, all our studied species are well separated (**Tables S2 and S3**). The similarity among all of the studied genomes of the *Salinispora* species showed clustering at an average nucleotide identity (ANI) (Figure 5), which follows the same sequence as the phylogenetic tree (Figure 4).

**Figure 4.**
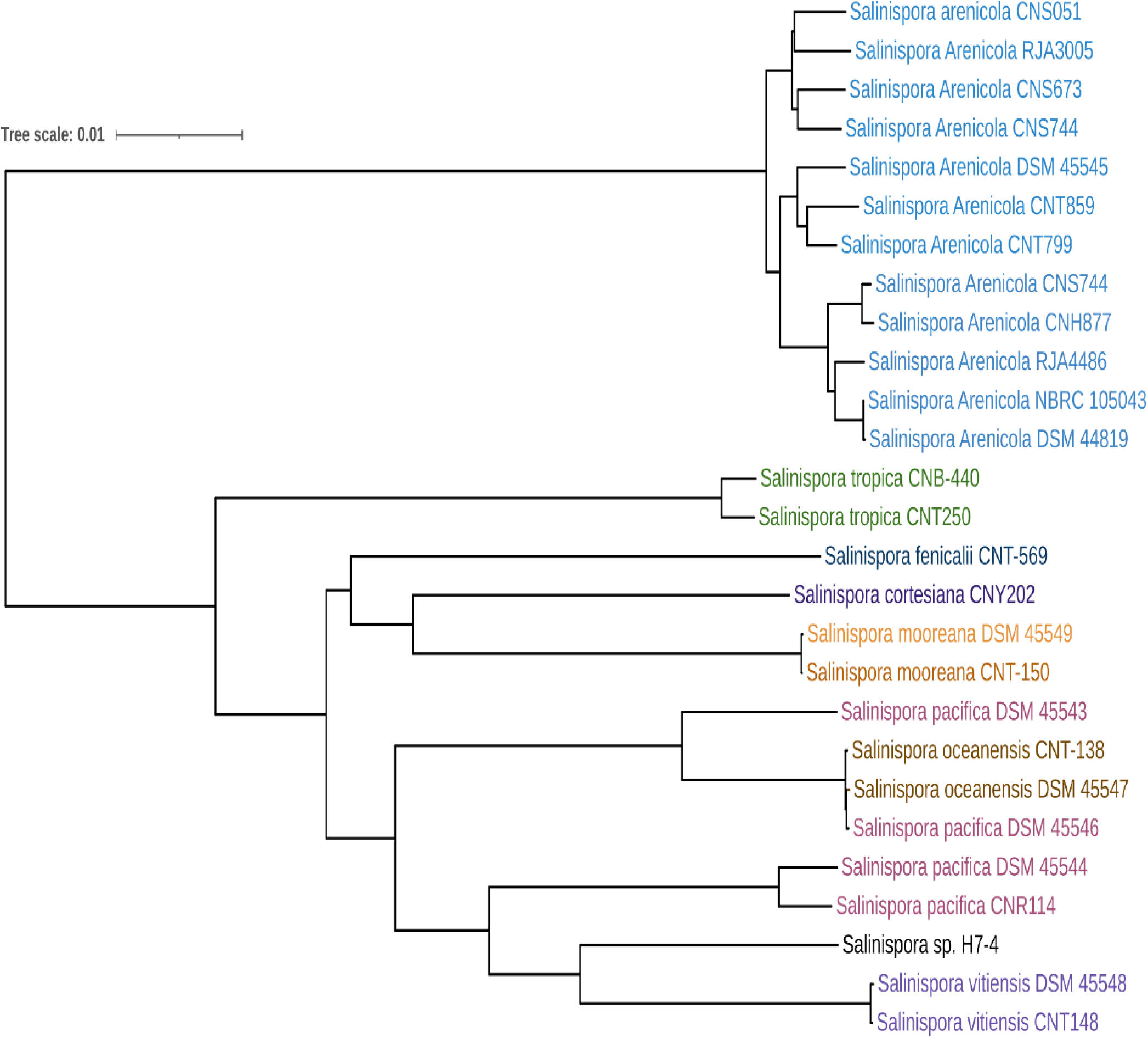
Tree inferred with FastME 2.1.6.1 (Lefort et al., 2015) from Genome BLAST Distance Phylogeny approach (GBDP) distances calculated from genome sequences. The branch lengths are scaled in terms of the GBDP distance formula, d_5_. The tree was rooted at the midpoint (Farris, 1972).

**Figure 5.**
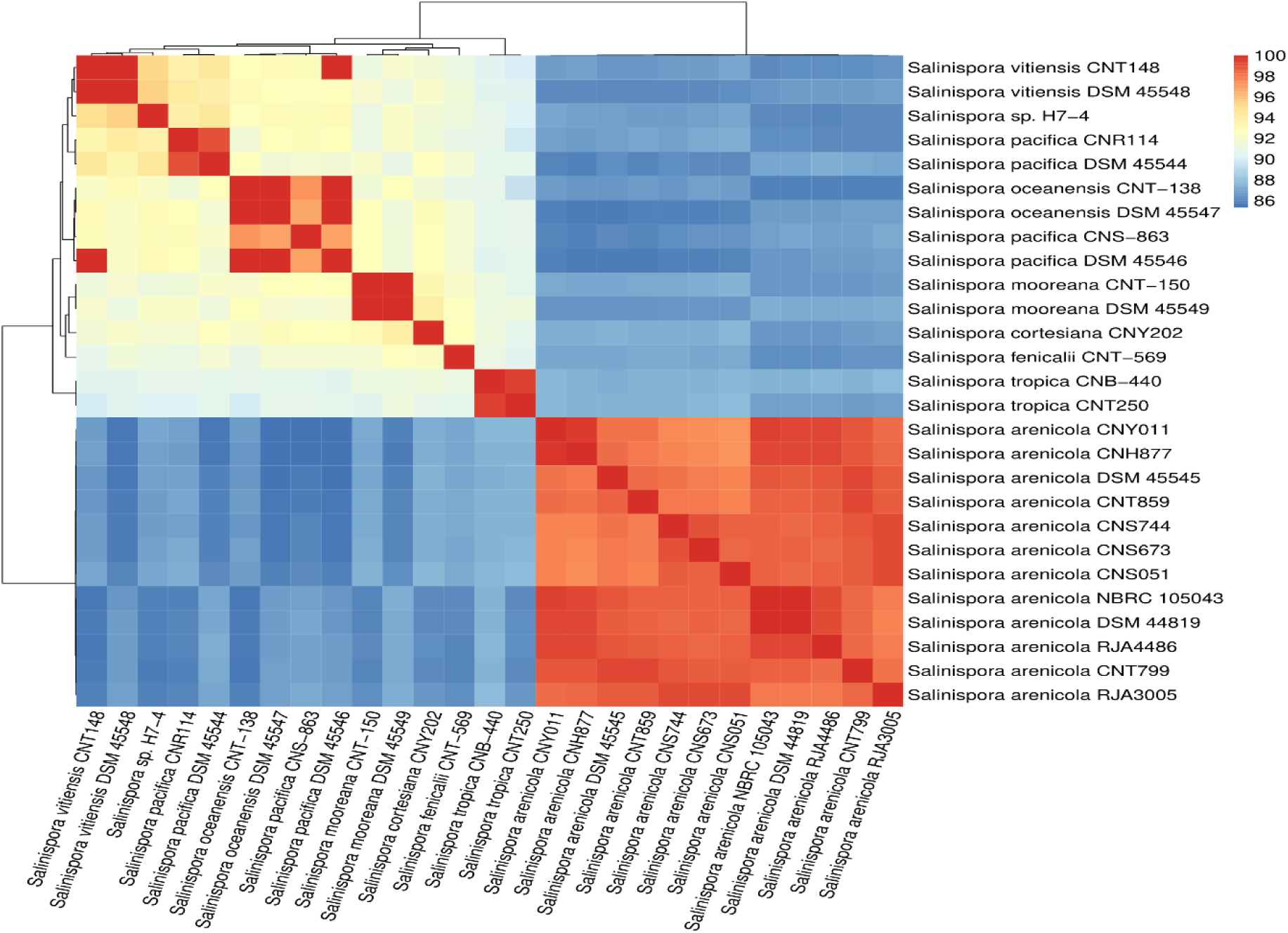
Cluster of the *Salinispora* genomes based on ANI values.

### Predictive Biosynthetic Gene Clusters (BGCs) of Salinispora species

When antiSMASH was used to look at the genomes of Salinispora, it was found that each genome has between 12 and 35 BGCs, with a mean of 26.07 and a standard deviation of 7.3. Among the total of 686 BGCs, the highest number of superclusters were present in both *Salinispora arenicola* CNH877 (Genome size 5.79 Mbp) and *Salinispora arenicola* CNT799 (Genome size 5.62 Mbp), while the lowest number were present in both *Salinispora vitiensis* DSM 45548 (Genome size 5.09 Mbp) and *Salinispora vitiensis* CNT148 (Genome size 5.06 Mbp). The smallest and largest genomes, respectively, featured 12 and 33 BGCs in *Salinispora mooreana* CNT-150 (Genome size 5.03 Mbp) and *Salinispora arenicola* CNS673 (Genome size 5.83 Mbp). The genome size was positively correlated with the number of BGCs per genome in a moderate but statistically significant way (Correlation coefficient, R = 0.648343932, coefficient of determination, R^2^ = 0.43618) (**Figure S4**). Gratifyingly, the number of predicted BGCs did not correlate with the number of contigs per genome (Correlation coefficient, R = 0.2756611; coefficient of determination, R^2^ = 0.075989), which gives a certain degree of robustness to our data, though actual BGC numbers might vary slightly; broken or merged clusters were not reinspected **(Figure S5)**.

In our study, we have found a variety of secondary metabolites producing genes that includes non-ribosomal peptides (NRPs), polyketides (PKs), ribosomally synthesized and post-translationally modified peptides (RiPPs), terpene, and PKS/NRPS hybrids. A total of 26 major BGC classes have been identified, such as polyketide synthases type I (T1PKS), non-ribosomal peptide synthetase (NRPS), terpene, polyketide synthases type II (T2PKS), NRPS-like, amglyccycl, N-acetylglutamine amide (NAGGN), polyketide synthases type III (T3PKS), butyrolactone, lanthipeptide class I, lanthipeptide class II, indole, ladderane, guanidinotides, NRPS independent siderophore, RiPP-like, blactam, phosphonate, thioamide-NRP, furan, LAP, other, RRE containing, transAT PKS, and hybrid, thoroughly scanning all *Salinispora* genomes (Figure 6).

**Figure 6.**
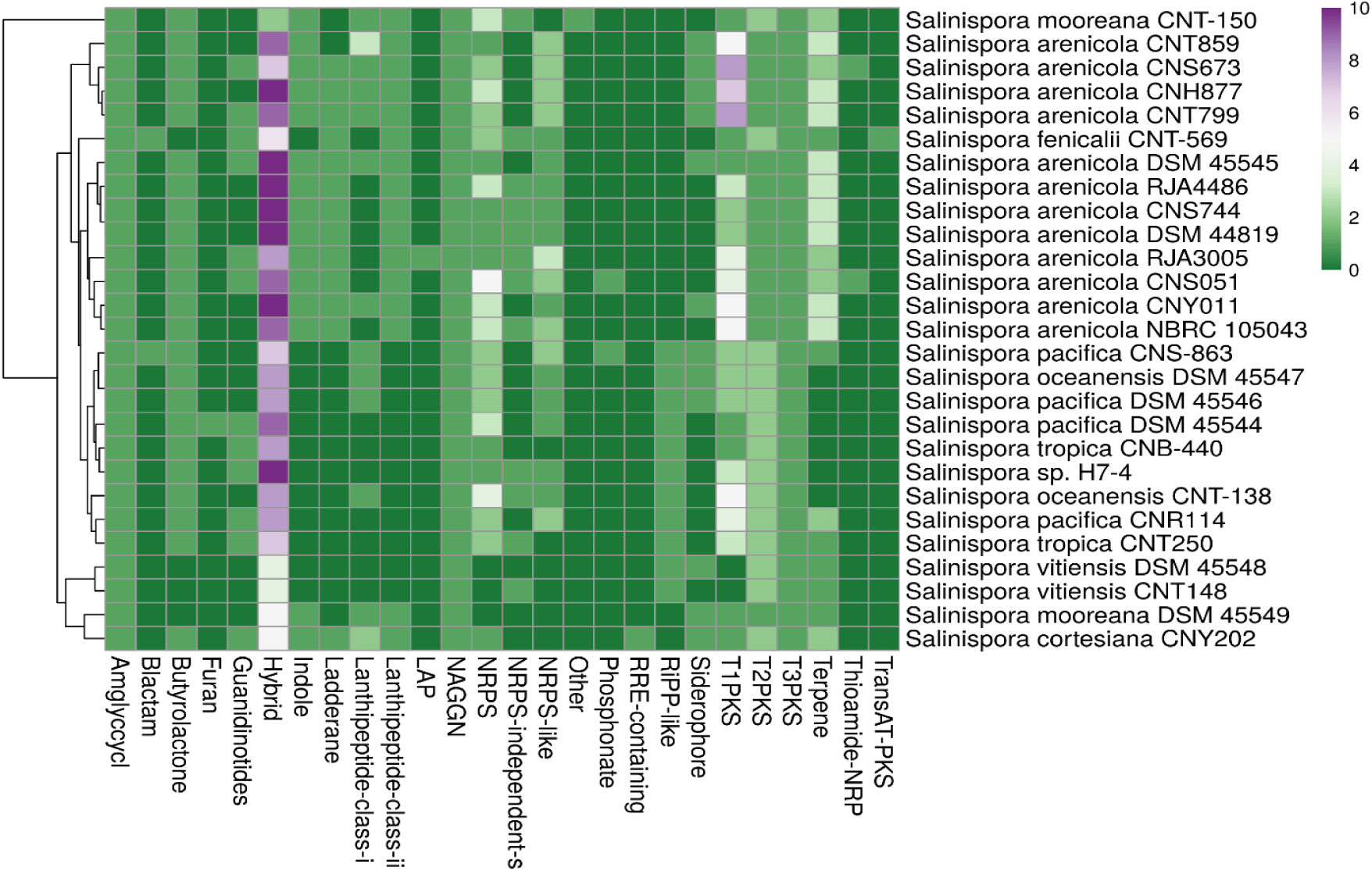
Heatmap of the antiSMASH database’s estimated distribution of BGC abundance in 27 *Salinispora* genomes.

Species of *Salinispora* are the repertoires of valuable components of biotechnological importance, as they carry 52 NRPS, 31 NRPS like BGCs, 84 T1PKS BGCs and 203 PKS/NRPS hybrid clusters with others. We have found diversified post-translationally modified peptides (RiPPs) such as 16 lanthipeptide-class I, 16 lanthipeptide-class II, 13 guanidinotides, 1 LAP, and 11 RiPP-like. Besides, 3 thioamide containing non-ribosomal peptides (thioamide-NRP), exceedingly rare BGCs have been identified. Biosynthetic Gene Clusters for common metabolites including 41 T2PKS, 27 T3PKS, 45 terpene, 27 amglyccycl, 27 NAGGN, 23 butyrolactone, 15 indole, 13 ladderane, 12 NRPS independent siderophores, 12 siderophore, 2 blactam, 2 phosphonate, 1 furan, 1 other, 1 RRE containing, and 1 transAT-PKS are also found.

Of the 27 *Salinispora* species, each contains at least two hybrid BGCs, and the highest number of hybrid BGCs (10) is found in *Salinispora arenicola* CNS673, *Salinispora arenicola* CNY011, *Salinispora arenicola* NBRC 105043, *Salinispora arenicola* DSM 44819, *Salinispora mooreana* CNT-150, *Salinispora tropica* CNB-440 and *Salinispora vitiensis* CNT148. Out of 210 hybrid BGCs, we have found 52 unique BGCs and NRPS-like/T1PKS clusters (61) as predominantly distributed hybrid clusters. According to our analysis, each T2PKS, amglyccycl, NAGGN and T3PKS BGCs has been found in all 27 *Salinispora* genomes, and T1PKS, NRPS, terpene, butyrolactone, and NRPS like BGCs have been found in 25, 24, 23, 23, and 20 *Salinispora* genomes, respectively (**Figure S6**)

These nine most common BGCs are responsible for 55.56% to 88.89% of all BGCs detected in a genome. In our study, we have found that a BGC class can be present in multiple copies in a strain. Out of 27 *Salinispora* genomes, 16 species contain multiple T1PKS BGCs, including *Salinispora arenicola* CNT799 and *Salinispora* sp. H7-4 with the highest copy number (8), and *Salinispora arenicola* NBRC 105043 with the second highest (7). Other BGCs that have multiple copies are NRPS, terpene, T2PKS, NRPS-like and lanthipeptide class I. Several BGCs have been detected in a small number of genomes, including transAT-PKS, furan, LAP, and RRE-containing in one genome, and blactam, phosphonate, and thioamide-NRP in only two genomes. These findings show that although many similar clusters exist in the genomes of different *Salinispora* species, the number may vary between species (Figure 7). Principal component analysis (PCA) based on BGCs data also demonstrated the consistent relationship among the *Salinispora* species, as *Salinispora mooreana* CNT-150 is found in separate positions from others (**Figure S7**).

**Figure 7.**
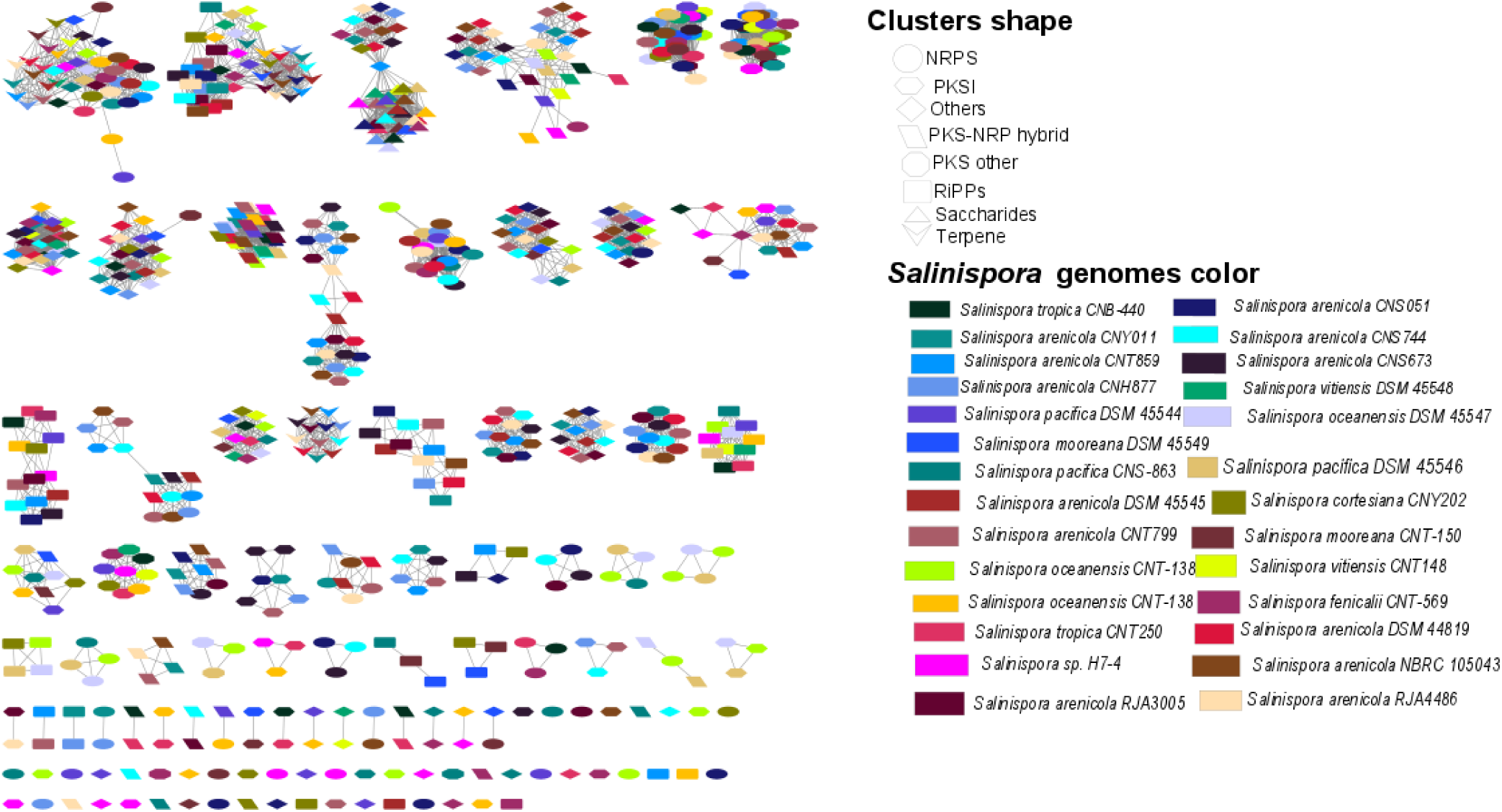
Sequence Similarity Networks (SSN) among the BGCs in *Salinispora* genomes.

A T1PKS cluster module may be complete or incomplete. When all keto synthase (KS), acyltransferase (AT) and acyl-carrier protein (ACP) domains are present in a module, it is complete, and in the absence of any of these essential domains, the module is incomplete. On the basis of the presence or absence of keto reductase (KR), dehydratase (DH) and enoyl reductase (ER) domains, complete modules are grouped into three categories: Reducing PKS module (RPKS modules based on the presence of two of KR, DH and ER domains), partially reducing PKS modules (PRPKS modules based on the presence of one of KR, DH and ER domains) and non-reducing PKS modules (NRPKS modules based on the absence of KR, DH and ER domains). All the analyses are based on the core biosynthetic genes of a cluster. The characterization of the PKS clusters at the structural level is important to identify compounds they may produce.

In our study, 686 BGCs were assigned to a total of 124 GCFs, indicating the promising diversity of BGCs in our dataset (**Table S4**). We have found 405 complete GCFs, and 281 fragmented GCFs, which maintained a positive and negative relationship with the number of contigs in a genome, respectively (**Table 2, Figures S8 & S9).** The relationship between each BGC and its respective GCF is determined by the calculation of distance (d), which value is classified into three categories: core (d≤900), putative (900 < d ≤ 1800) and orphan (d >1800) (Hou et al., 2023; Kautsar, Van Der Hooft, et al., 2021). In our findings, 454 (66.2%) BGCs were “core” GCF, which indicated that those BGCs were relatively conserved; 225 (32.8%) BGCs were “putative” GCF, implying that they were potential candidates for encoding previously unknown compounds, and 7 (1%) BGCs were subclassified as “orphan”, which were the best candidates for producing highly novel secondary metabolites as they were highly distant from the known BGCs. The orphan BGCs belonged to T1PKS-arylpolyene-other-NRPS hybrid (1), T2PKS-lanthipeptide hybrid (1), NRPS-T1PKS hybrid (2), other-NRPS like hybrid (1), and NRPs (2). The putative BGCs were mostly belonged T1PKS (43) followed by NAGGN (27), NRPS (26), T1PKS-NRPS like hybrid (23), guanidinotides(14), T2PKS (13), siderophore (12), lanthipeptide (11), NRPS like (9), NRPS-T1PKS hybrid (7), NRPS-arylpolyene-T1PKS-ladderane hybrid (4), NRPS-T1PKS-NRPS like hybrid (3), T1PKS-RRE containing hybrid (3), T3PKS (3), NRPS -NRPS like hybrid (2), NRPS-arylpolyene hybrid (2), NRPS -lanthipeptide hybrid (2), indole (2), LAP-thiopeptide hybrid (2) NRPS-ladderane hybrid (1), NRPS-other-oligosaccharide hybrid (1), NRPS-PKS like hybrid (1), NRPS-T1PKS-Terpene hybrid (1), NRPS-T1PKS-Terpene-LAP-thiopeptide hybrid (1), thioamide-NRP (1), blactam (1), furan (1), other (1), other-T1PKS hybrid (1), phosphonate (1), T1PKS-NRPS like-transAT PKS like hybrid (1), T1PKS-oligosaccharide-NRPS like-PKS like hybrid (1), LAP-siderophore-thiopeptide (1), RRE containing (1), amglyccycl-ladderane hybrid (1), and Terpene-LAP-thiopeptide hybrid (1).

**Table 2.**
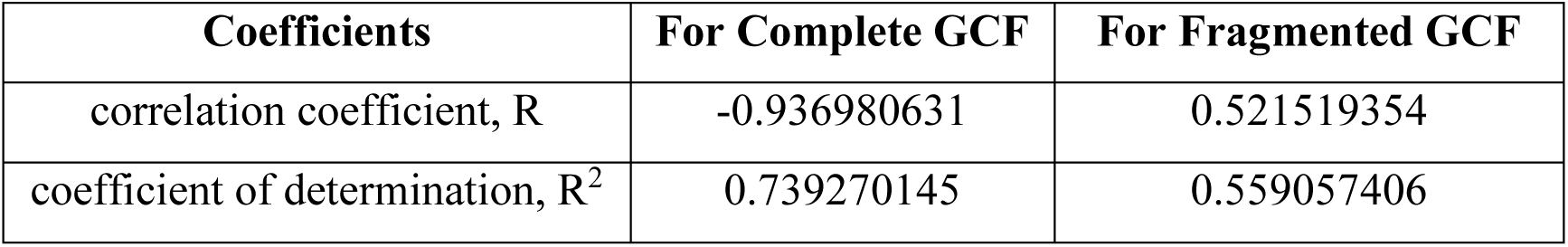
Relationship of complete or fragmented GCF and number of contigs in a genome.

BiG-SCAPE was used to make a map of how the genomes of all BGCs predicted by antiSMASH are put together. This tool creates a sequence similarity network (SSN) and groups similar BGCs into gene cluster families (GCFs) to map their diversity and evolution. The generated SSN clearly confirms the diversity of the predicted BGCs, as indicated by 148 distinct GCFs, 51 of which are singletons (Figure 7).

Out of 686, 553 BGCs were related to various known compounds, and 133 BGCs were functionally unknown, which indicates the potentiality of *Salinispora* species as sources of secondary metabolites. Gene clusters of *Salinispora* species compared with known clusters in the antiSMASH database were antimicrobial, antibiotic, antibacterial, antifungal, anti-cancer, anti-tumor, antioxidant, immunosuppressive, etc. (**Figure S10 and Table S5**).

From the BAGEL4 data analysis, we identified 89 bacteriocin coding clusters for the whole genome datasets of *Salinispora* species (**Table S6**). Most of the bacteriocins found here were annotated as class II bacteriocins which are antimicrobial peptides and have not undergone posttranslational modification.

### KS and C Domain Determination in the Salinispora Genomes Using NaPDoS2

The ketosynthase (KS) and condensation (C) domains, respectively, represent the existence of BGCs for PKs and NRPs. We have found a total of 684 KS domains (99.71%) and 531 C domains (77.41%) among the total of 686 BGCs predicted from the 27 *Salinispora* reference genomic sequences (**Table S7**). The most KS domains (39) were found in *Salinispora arenicola* DSM45545 and the least KS domains (12) were seen in the *Salinispora mooreana* CNT-150 genome while the average number of KS domains was found to be 25.33. On the other hand, the highest number of C domains (30) were found in *Salinispora fenicalii* CNT-569, and *Salinispora arenicola* CNS051. *Salinispora arenicola* DSM448194 and *Salinispora arenicola* RJA3005 both had the fewest C domains (8), and the overall average C domain count was 19.67.

The total of 684 KS domains present in each of the *Salinispora* genomes were distributed in eight classes; among them, most (375) were from the modular class, and the rest were iterative (62), metazoan (1), trans (9), aromatic (88), FAS (78), polyene (68) and unclassified (3) (**Figure S11)**. Iterative PKS employs the same domain numerous times, whereas modular PKS enzymes are big multi-domain enzymes that only use each domain once during the synthesis process [20]. The similarity among KS domains of all strains ranged between 33 and 100%, with an average similarity of 69.72%.

The sequence of this class were similar to genes associated with various biosynthetic pathways, which lead to the formation of rifamycin (108), xantholipin (27), vicenistatin (1), urdamycin (1), tylactone (1), tubulysin (5), totopotensamide (1), tiacumicin (12), tartrolon (1), FAS (73), streptolydigin (4), sporolide (2), spinosyn (2), sorangicin (2), skyllamycin (14), simocyclinone (1), sceliphrolactam (1), sanglifehrin (2), salinosporamide (8), salinomycin (1), salinilactam (37), rubromycin (1), rhizoxin (1), reveromycin (6), pyrronazol (1), patellazole (1), padanamide (20), pactamycin (2), oxazolomycin (1), oviedomycin (1), oligomycin (9), nostophycin (2), nostopeptolide (5), neocarzinostatin (4), naphthomycin (4), nanchangmycin (1), myxovirescin (1), myxothiazol (1), mycocerosic acid (1), meridamycin (13), megalomicin (5), maduropeptin (1), lymphostin (24), lomaiviticin (18), leinamycin (1), herboxidiene (9), halstoctacosanolide (4), fostriecin (9), fasamycin (28), erythromycin (4), epothilone (8), enterocin (4), cyanosporaside (21), curacin (13), colabomycin (18), calicheamicin (25), avermectin (2), asukamycin (36), and apoptolidin (3).

The total of 531 C domains present in each of the *Salinispora* genomes were distributed in seven classes; among them, the most abundant (362) were from the LCL class, and the rest were condensation (1), cyclization (108), DCL (26), epimerization (6), modified amino acids (26), and starter (2) (**Figure S12)**. The LCL domains were found in all species, and the condensation domain was found only in *Salinispora oceanensis* CNT-138. The LCL domain catalyzes the formation of a peptide bond between two L-amino acids, while the DCL domain catalyzes the formation of a peptide bond between an L-amino acid and a growing peptide ending with a D-amino acid. Starter domains acylate the first amino acid with a fatty acid, polyketide, or other molecule, and the cyclization domain catalyzes both peptide bond formation and the subsequent cyclization of cysteine, serine or threonine residues. The epimerization domain changes the chirality of the last amino acid in the chain from L to D; modified AA modifies the incorporated amino acid; and the condensation domain’s functionality is unknown [21]. The seven classes of C domains were involved in the biosynthesis of nostopeptolide (145), actinomycin (13), anabaenopeptilide (66), pyoverdine (23), pyridomycin (12), arthrofactin (12), syringomycin (15), bacitracin (5), balhimycin (7), bleomycin (70), complestatin (3), cyclomarin (8), microcystin (28), pristinamycin (24), sporolide (29), thiocoraline (12), tubulysin (33), tyrocidine (1), and yersiniabactin (10). The similarity among C domains of all strains ranged between 23 and 100%, with an average similarity of 39.6%, which is very low and indicates their natural product diversity.

Although the same product forming pathways were found in multiple species, phylogenetic analysis of NRPS C domains and PKS KS domains clustered them differently based on their sequence similarity (Figures 8 **& 9**). If the sequence similarity with the biosynthetic domains of experimentally validated compounds is > 90%, an accurate structural class of the compounds can be anticipated from these domains [21]. However, this research showed that these domains did not meet the aforementioned requirements, making it obvious that they may serve as a repository for novel biomolecules.

**Figure 8.**
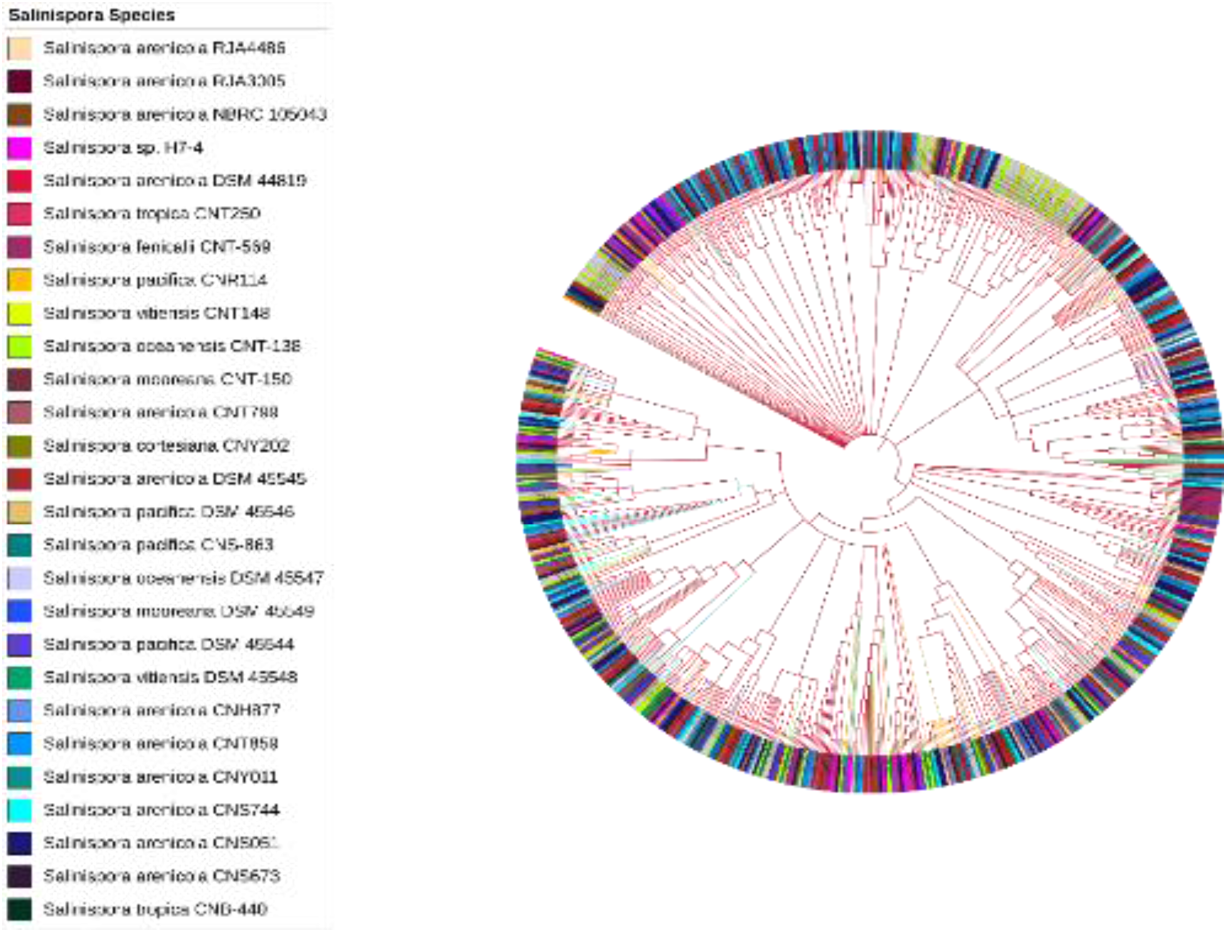
Phylogenetic analysis of ketosynthase (KS) domains of PKS genes in *Salinispora* genomes. Different *Salinispora* species are labeled in colors, while colored branches display different KS domains (Crimson= Type I modular cis-AT, Spring Bud= Type I trans-AT, Yellow= Type II aromatic, Purple= Type II FAS, Green= Type II polyene, Cyan= Type I iterative cis-AT, Blue= Type I Metazoan-type PKS).

**Figure 9.**
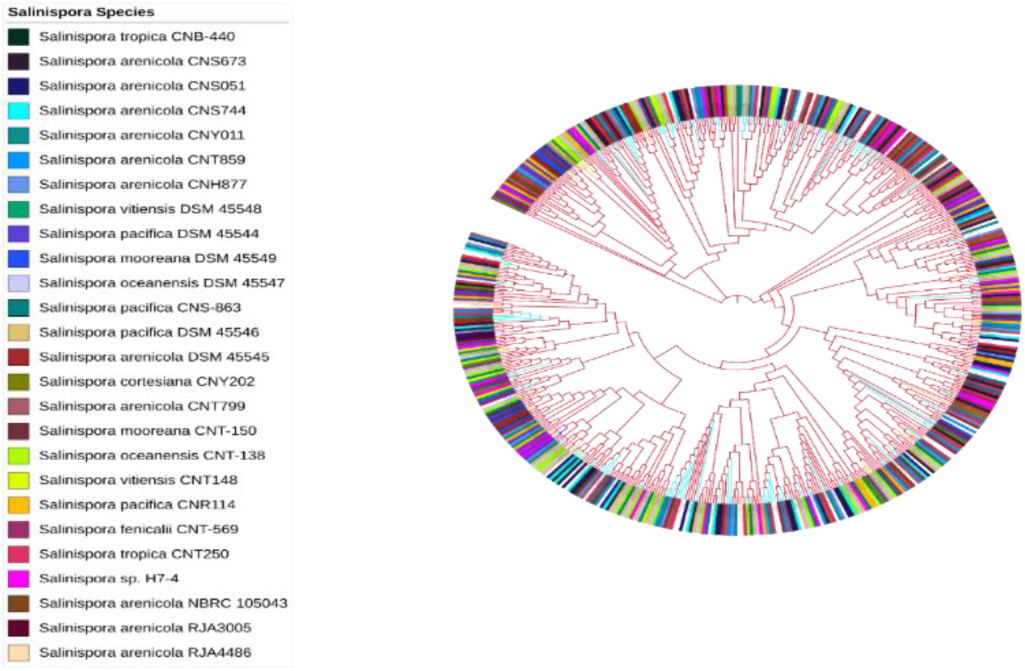
Phylogenetic analysis of condensation (C) domains of NRPS genes in *Salinispora* genomes. Different *Salinispora* species are labeled in colors, while colored branches display different C domains (Crimson= LCL, Spring Bud= epimerization, Yellow= modified amino acid, Purple= DCL, Green= Condensation, Cyan= cyclization, Blue= starter).

### Abundance of CAZY Enzymes

In our studied *Salinispora* species, a total of 3604 genes were involved in the production of CAZY enzymes, which are organized into six families: families of auxiliary activities (AA), carbohydrate-binding modules (CBM), carbohydrate esterases (CE), glycoside hydrolases (GH), glycosyl transferases (GT), and polysaccharide lyases (PL) (**Table S8**). The majority of the enzymes in these families came from GT (1548), followed by GH (898) and CE (251), with PL enzymes (4), which were detected in *Salinispora vitiensis* DSM 45548, *Salinispora cortesiana* CNY202, *Salinispora vitiensis* CNT148, and *Salinispora sp.* H7-4, providing the lowest number of enzymes. *Salinispora pacifica* DSM 45546 and *Salinispora oceanensis* CNT-138 represented the highest number of CAZY enzymes (145), whereas *Salinispora mooreana* CNT-150 represented the lowest number of CAZY enzymes (109). A total of 62 various categories of the major six CAZY enzyme families were found in our study and among them, AA4, CBM11, GH12, GH20, GH38, GH48, GH6, GH65, GH85, GT20, GT26, GT28, GT39, and GT51 (14 families) were present in all species with the same copy number (Figure 10). Moreover, the CAZyme gene cluster (CGC) finder predicted 1665 clusters in the 27 *Salinispora* genomes, and 61.67 was the average. Some enzymes were predicted to be secreted freely in the presence of a signal peptide, and some were thought to be anchored to the cell in the absence of any predicted signal peptide (**Dataset S1**). Similarity in CAZymes was shown in PCA, which was consistent with their cladogram (**Figure S13**).

**Figure 10.**
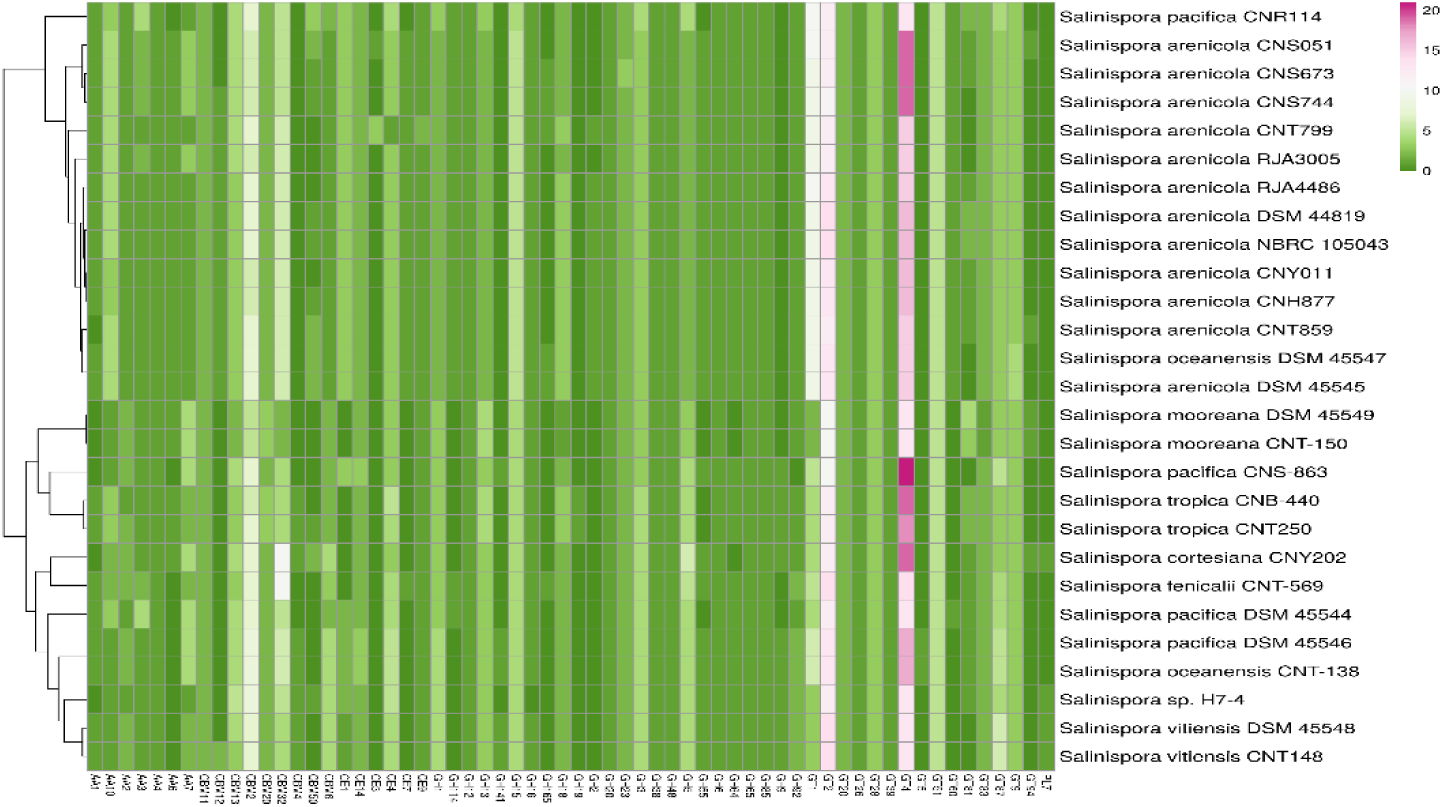
A clustered heatmap represents the abundance of different categories of CAZY enzymes in *Salinispora* genomes.

## Discussion

Bioactive natural chemicals and their secondary metabolites, which are crucial for both biotechnology and medicine, have been the main source of new therapeutic agents. Additionally, in the genomic era, when the number of available genomes is increasing, genome mining for the finding of cryptic biosynthetic areas, together with synthetic biology, is incredibly helpful in drug development. The main goal of this research was to use a variety of bioinformatics techniques and all the publicly available NCBI Refseq genome data for the *Salinispora* species to identify genetic clusters responsible for putative drug-like metabolites. Our results showed that *Salinispora* species have a significant and distinct natural product metabolic potential with high diversity, despite earlier comprehensive studies, demonstrating that they are still a good source of novel metabolites [22]. In our study we selected only NCBI Reference Sequence (RefSeq) data as it provides a comprehensive, integrated, non-redundant, well-annotated set of sequences. We have found the most abundant COGs proteins were related to transcription, amino acid transport and metabolism, coenzyme transport and metabolism, translation, ribosomal structure and biogenesis, and lipid transport and metabolism in the species of *Salinispora*. Except for *Salinispora oceanensis* DSM 45547 and *Salinispora pacifica* DSM 45546, all species were clearly distinct from one another according to genome-based comparison using the ANI and dDDH values. These species’ branch lengths were identical according to the phylogenomic analysis, which further suggests that they might be the same species. Core protein-based phylogenetic analysis of the aforementioned species as well as WGS-based phylogenetic analysis both supported the same conclusion. The purpose of this work was to gain knowledge about the genes that produce bioactive and CAZY enzymes in the 27 *Salinispora* species of actinobacteria. We have found that the genome sizes of the *Salinispora* species varied and that the number of BGCs and the genome sizes correlated well. According to BiG-SCAPE’s analysis, the top order of detected BGCs included 194 others (amglyccycl, indole, CDPS, furan, butyrolactone, etc.), 97 PKSI, 95 NRPS, 81 PKS others, 75 Hybrids (PKS-NRPS), 74 RiPPs, 44 Terpene, and 26 Saccharides. At least one cluster of T2PKS, amglyccycl, NAGGN, and T3PKS was present in every genome. In comparison to all other species, *Salinispora arenicola* CNH877 displayed the most frequent clusters of T1PKS, NRPS, NRPS-like, and PKS/NRPS hybrids but *Salinispora arenicola* RJA3005 and *Salinispora oceanensis* DSM 45547 showed no clusters of T1PKS, NRPS, or NRPS-like.

In addition to the majority of NRPSs, NAGGN BGCs common in *Salinispora* predicted for new compounds, *Salinispora* genomes carry some BGCs for NRPS-like type compounds, similar to stenothricin, rifamycin, legonindolizidine A6, RiPP-like type compounds, similar to lipstatin, saframycin A/saframycin B, T2PKS type compounds, similar to lomaiviticin A/lomaiviticin C/lomaiviticin D/lomaiviticin E, paramagnetoquinone 1/paramagnetoquinone 2, formicamycins A-M, pradimicin-A, T3PKS type compounds, similar to loseolamycin A1/loseolamycin A2, terpene type compounds, similar to carotenoid, isorenieratene, rifamycin, frankiamicin, indole type compounds, similar to staurosporine, ladderane type compounds, similar to triacsin C, and NI-siderophore type compounds, similar to FW0622. Additionally, unidentified BGCs have been discovered for NAGGN, amglyccycl, lanthipeptide class I, lanthipeptide class II, LAP, NRPS, NRPS-like, phosphonate, RiPP-like, RRE-containing, T1PKS, and terpene, which seem to be cryptic gene clusters [23]. We have identified very comparable clusters (> 50%) in the *Salinispora* genome that resemble well-known chemicals that have been chemically described and are used for a variety of applications. These products are benzastatin J, enterocin, fredericamycin A, FW0622, ketomemicin B3/ketomemicin B4, lomaiviticin A/lomaiviticin C/lomaiviticin D/lomaiviticin E, loseolamycin A1/loseolamycin A2, lymphostin/neolymphostinol B/lymphostinol/neolymphostin B, naphthyridinomycin, peucechelin, polyoxypeptin, prejadomycin/rabelomycin/gaudimycin C/gaudimycin D/UWM6/gaudimycin A, rifamycin, salinilactam, salinipostin G, salinosporamide A, sporolide A/sporolide B, staurosporine, thiolactomycin, triacsin C, and valclavam/(-)-2-(2-hydroxyethyl)clavam. Furthermore, the *Salinispora* BGC clusters with less than 50% sequence similarity represent a possible natural product repertoire with a broad range. In the identified compounds with high and low sequence similarities, antimicrobial, antibiotic, antibacterial, antifungal, anticancer, anti-tumor, antioxidant, and immunosuppressive activities were found. According to what we know, this genus of actinobacteria was not extensively employed to explore the aforementioned bioactive substances; as a result, additional research will be required to extract the compounds, characterize them, or activate cryptic genes in vitro if that doesn’t work. In addition to producing bioactive substances, these bacteria are useful in the agricultural and environmental fields because they produce siderophores and have enzymes (AA, CBM, CE, GH, GT, and PL) that are active on carbohydrates [23–28].

## Conclusions

The continuous resurgence in natural product research has been fueled by the development of new technologies in genome sequencing, bioinformatics, molecular genetics, and a better understanding of the metabolic principles that control natural product assembly. By looking at the *Salinispora* genome, we have found a number of secondary metabolites and carbohydrate-active enzymes, which could help develop and find new drugs to fight new pathogens. We discussed the genomic characteristics, phylogenomic, core-proteomics based phylogenetic analysis, numerous secondary metabolites generating genes and their potential products, networking among BGCs, polymer degrading enzymes, and genomic properties of all *Salinispora* species. With the wide variety of BGCs, this rare species of actinobacteria has the potential to produce distinctive putative products, and more research into these prospective secondary metabolite coding clusters (BGCs) will probably produce more relevant information. Target-specific products that are only present in specific species may be made available with the aid of this research. *Salinispora* may also be employed in the agricultural industry since it can degrade a variety of polymers and because the crop plant satisfies a range of nutritional needs. Further investigation of the potential secondary metabolite coding clusters (BGCs) present in *Salinispora* genomes will likely yield more useful information.

